# Identifying key factors for improving ICA-based decomposition of EEG data in mobile and stationary experiments

**DOI:** 10.1101/2020.06.02.129213

**Authors:** Marius Klug, Klaus Gramann

## Abstract

Recent developments in EEG hardware and analyses approaches allow for recordings in both stationary and mobile settings. Irrespective of the experimental setting, EEG recordings are contaminated with noise that has to be removed before the data can be functionally interpreted. Independent component analysis (ICA) is a commonly used tool to remove artifacts such as eye movement, muscle activity, and external noise from the data and to analyze activity on the level of EEG effective brain sources. While the effectiveness of filtering the data as one key preprocessing step to improve the decomposition has been investigated previously, no study thus far compared the different requirements of mobile and stationary experiments regarding the preprocessing for ICA decomposition. We thus evaluated how movement in EEG experiments, the number of channels, and the high-pass filter cutoff during preprocessing influence the ICA decomposition. We found that for commonly used settings (stationary experiment, 64 channels, 0.5 Hz filter), the ICA results are acceptable. However, high-pass filters of up to 2 Hz cutoff frequency should be used in mobile experiments, and more channels require a higher filter to reach an optimal decomposition. Fewer brain ICs were found in mobile experiments, but cleaning the data with ICA proved to be important and functional even with 16 channels. Based on the results, we provide guidelines for different experimental settings that improve the ICA decomposition.

## 1 INTRODUCTION

Over the last decade, the development of lightweight portable electroencephalography (EEG) amplifiers and new data-driven analyses approaches led to the investigation of the neural basis of ecologically valid cognitive processes in actively behaving human participants outside established laboratory environments. Experiments now allow active behavior of participants both in the lab (e.g. Gramann et al., 2010; De Sanctis et al., 2014; Ehinger et al., 2014; Gehrke et al., 2018; Djebbara et al., 2019) and in the real world, which increases our understanding of human brain dynamics accompanying embodied cognitive processes as well as the impact of real world environments (e.g. Debener et al., 2012; Wascher et al., 2014; Ladouce et al., 2017; Protzak and Gramann, 2018; Wunderlich and Gramann, 2018). While these experimental protocols provide new insights into the neural activity subserving cognition in more realistic and natural settings, they present new challenges. Mobile EEG or Mobile Brain/Body Imaging (MoBI; Makeig et al., 2009; Gramann et al., 2011, 2014; Jungnickel et al., 2018) recordings are impacted by the participant movement due to movement-related electrical activity stemming from facial muscles, neck muscles and eye movements that naturally accompany active behaviors. While these physiological sources are usually unwanted but may still be regarded as providing additional insights into the ongoing cognitive processes, other artifactual contributions to the recording are even less welcome. For example, movement in mobile protocols might lead to cable sway or micro movement of electrodes that contribute artifactual activity into the recording. Finally, environmental sources and the equipment necessary for the experiment itself like head mounted virtual reality (VR) systems or treadmills can be another unavoidable source of artifacts in mobile recordings. Due to volume conduction, all these signals mix at the sensor level and render it difficult to dissociate brain from non-brain activity to investigate the neural basis of the cognitive processes of interest. While movement-related non-brain sources are specifically problematic for experiments including actively behaving participants, contributions from sources like eye movements, facial muscles, and neck muscle activity can also be found in EEG data recorded in established desktop settings. These forms of biological signals are traditionally considered artifacts and are one of the main reasons why established EEG research minimizes any kind of participant movement, including eye movements or blinks. Thus, the ability to interpret EEG data from both classic stationary as well as MoBI experiments depends greatly on the ability to dissociate signals of interest originating in the brain from those of other sources.

This dissociation can be achieved by applying spatial filter methods to the data, exploiting the fact that electrical activity is recorded with multiple electrodes on the scalp. Among different spatial filter approaches, blind source separation techniques (Bell and Sejnowski, 1995; Makeig et al., 1997; Hyvärinen et al., 2001) proved to be very successful and specifically independent component analysis (ICA) applied to EEG and magnetoencephalography (MEG) data demonstrated increasing popularity among researchers. While the number of ICA applications to EEG data constantly grew over the last 25 years from 16 papers in 1995 to 5450 papers in 2019 alone (search term “EEG” + “Independent component analysis”, Google Scholar, accessed on 2020-05-18), the variations of preprocessing the data to eventually applying ICA also increased. In most cases a channel density of 64 and upwards is being used for ICA since spatial filtering typically improves with more degrees of freedom, but less consensus is reached considering the applied filter. Often a high-pass filter of 1 Hz is used, but lower frequencies like 0.5 Hz or higher ones like 2 Hz or even 3 Hz can be found in the literature as well. Sometimes, additional low-pass filters are applied while many times none is used. While some of the preprocessing steps have been evaluated regarding their impact on the subsequent ICA decomposition, not all factors have been systematically investigated yet. The purpose of this study is to shed light on the relevant contributions of different factors on ICA for both stationary and mobile experiments, and to provide a “best practices” guideline to improve the ICA decomposition.

In this paper, we will first discuss the underlying assumptions of EEG source separation, as well as prior research on the effect of different data preprocessing settings on ICA. Formulating our hypotheses, we will then present our approach to investigate the impact of the three most common factors influencing ICA decompositions: high-pass filter settings, channel density, and movement. Finally, the results will be discussed and recommendations will be given.

## 2 THE EEG LINEAR MIXING MODEL

Analyzing EEG data with ICA is based on a general assumption that the data matrix *X* ∈ ℝ^*N*^ × *M* recorded by the EEG electrodes is a linear mixture of different sources *S* ∈ ℝ^*N*^ × *M* with a mixing matrix *A* ∈ ℝ^*N* ×*N*^ :

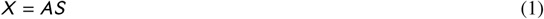

with *N* being both the number of sources and EEG channels, and *M* being the number of samples in the dataset (Hyvärinen et al., 2001; Hyvärinen and Oja, 2013). Without loss of generality, sources are assumed to be temporally and spatially stable, and statistically independent with unit variance. These assumptions can now be leveraged to compute an inverse un-mixing matrix *W* = *A*^−1^ (*W* ∈ ℝ^*N* ×*N*^), such that

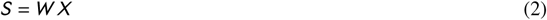

Since the model is linear, equation 2 holds for individual independent components (ICs) as well, which can then be examined on their topography and other evaluations to determine whether or not they are considered artifactual and should be removed. Cleaning the data by removing an independent component is achieved by simply removing both the respective column of *S* and the associated row of *A*, which can then be multiplied again to generate a cleaned dataset:

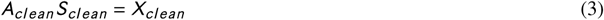

Finding *W* is an ill-posed problem without an analytical solution. Different ICA algorithms use different heuristcs and thus compute slightly different un-mixing matrices (Hyvärinen et al., 2001), and even the same algorithm does not always converge on the same solution for the same data (Groppe et al., 2009; Artoni et al., 2014).

### 2.1 Achieving an Optimal Decomposition

Since ICA is becoming increasingly popular for EEG research, efforts have been made to identify the best algorithms and prerequisites to obtain a good decomposition of the data. Comparing different algorithms, Delorme et al. (2012) and Leutheuser et al. (2013) found that AMICA (Palmer et al., 2011) performed best among different algorithms. This was confirmed in part by Zakeri et al. (2014), but it was concluded that the choice of preprocessing was more relevant to the decomposition quality than the algorithm itself.

Already early work on ICA has found this to be relevant, as “[t]he success of ICA for a given data set may depend crucially on performing some application-dependent preprocessing steps” (Hyvärinen et al., 2001, p. 263). One often used method to improve the decomposition quality is that of high-pass filtering. High-pass filtering is essentially another linear transformation of the data and thus does not violate the ICA assumptions, as it can be expressed by multiplying equation 1 with a filtering matrix *F* from the right:

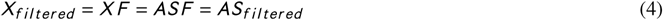

Since the un-mixing matrix *A* remains unchanged and only the reconstructed sources are filtered with the same filter it is possible to compute *A* on filtered data and apply it to the unfiltered data (Hyvärinen et al., 2001), which is common practice in ICA analysis. While low-pass as well as high-pass filtering may remove noise from the data, filtering also bears the risk of removing relevant information. For low-pass filtering this is the case especially for high-frequency activity stemming from muscle contractions while for high-pass filtering this concerns slow cortical potentials in the data. Noise in form of slow drifts in the data often occurs in all or many EEG channels and is thus hard to separate (Winkler et al., 2015). Removing slow drifts can thus benefit the decomposition. While being used in practice almost universally as a preprocessing step before ICA, the exact filter specifications, especially the cutoff frequency of the high-pass filter, are not always agreed on.

Several studies have investigated the effect of high-pass filtering on the ICA decomposition. Groppe et al. (2009) have found that removing the mean of epoched data (which acts as a leaky high-pass filter) improved the decomposition greatly. Following up on this result, (Zakeri et al., 2014) compared the effects of filtering, epoching, de-meaning, and including electrooculography and electrocardiography channels (EXG) on the ICA decomposition. They found that the best approach was to compute the ICA on filtered continuous data including EXG. However, the applied filter was a band-pass filter of 0.16-40 Hz, which is a low high-pass filter compared to previous studies that used filters of 0.5 Hz or higher (Delorme et al., 2012; Leutheuser et al., 2013). Additionally, the application of a band-pass filter does not allow any conclusions for high-pass filtering specifically. This was addressed in detail by Winkler et al. (2015), who compared the effect of high-pass filtering in frequencies of 0 (no filter) to 40 Hz on the ICA decomposition. It was found that filters of <0.5 Hz were indeed suboptimal, and the best results were achieved with filters of 1-2 Hz. In a recent study, Frølich and Dowding (2018) found that filtering data that had already been band-pass filtered from 2-40 Hz with another 14 Hz high-pass filter led to better decompositions in a scenario with high muscular contributions to the data. Considering especially the impact on data with high amounts of ocular movements, Dimigen (2020) found that a high-pass filter cutoff of 1-1.5 Hz produced best results when comparing filters of 0 (no filter) to 30 Hz. In addition, the study specifically investigated low-pass filtering and found that low-pass filtering with 40 Hz was detrimental compared to 100 Hz. Lastly, in a study using a phantom head to simulate EEG recording during walking, Richer et al. (2019) found that adding EMG channels to the recording improved the cleaning capabilities of ICA.

Taken together, previous studies suggest that a high-pass filter between 1 and 2 Hz and no low-pass filter seems to be the best choice and may improve the decomposition. However, the results are inconclusive, and two factors have not yet been addressed that can be observed in several EEG studies. Firstly, no study yet compared the different requirements of standard stationary experiments and active MoBI experiments. While (Winkler et al., 2015) used a stationary experimental protocol where comparatively low amounts of artifacts were to be expected, other studies only investigated muscle artifacts (Zakeri et al., 2014; Frølich and Dowding, 2018; Richer et al., 2019), with one study exploring the removal of ocular artifacts in great detail (Dimigen, 2020). The second not yet examined factor is the number of channels which were used for the decomposition, as none of the above-mentioned studies compared scalp channel montages, and the reported studies used channel densities ranging from 45 to 71 channels. Yet, as the number of cortical and artifactual sources stays the same, no matter how many channels are used, the distinction between signal and noise could become more evident with an increasing number of channels, as the sources might be more clearly separated. This is especially important in mobile studies as more contributions from physiological sources (eye and muscle activity) and other artifacts (cable sway, electrical noise) are expected in these types of experiments. Here, the available degrees of freedom for the decomposition might play a crucial role.

## 3 CURRENT STUDY

We thus specifically asked how movement in EEG experiments would influence the quality of the decomposition. We were further interested whether and how the number of channels would impact the decomposition results. Lastly, we investigated how the filter settings during preprocessing influence the outcome of the decomposition, especially considering the differences between stationary and mobile experiments with different spatial densities of the montages. The quality of the decomposition was assessed using the dipolarity of IC topographies, the noise in the event-related potential data after backprojection to the sensor level using only brain ICs, and the number of brain components automatically classified from all resultant ICs. We assumed that higher-density channel montages would be beneficial in general, and more so for data of mobile experiments. Especially in the mobile setting we expected that adding EMG data of the neck would improve the decomposition. We further expected the best decomposition results for preprocessing with a high-pass filter cutoff in the range between 0.5-2 Hz. Finally, we hypothesized that mobile experiments include more slow drifting signals due to mechanical and movement-related artifacts and thus require a higher cutoff frequency than stationary experiments to achieve the best decomposition.

## 4 METHODS

We analyzed data from a spatial orientation MoBI experiment, which is particularly useful for this study as it included a stationary as well as a mobile condition in which participants solved the identical task with comparable visual input. It allows for a comparison of stationary and mobile EEG setups and thus the impact of movement on decomposition quality. Details of the experiment can be found in Gramann et al. (2018).

### 4.1 Experiment and Dataset

#### 4.1.1 Participants

20 healthy adults participated in the study (11 females, aged 20-46 years, M=30.25 years) and were compensated with either 10€ /h or course credits. One participant aborted the experiment due to motion sickness, the remaining 19 datasets were used for analysis. The experiment was approved by the local ethics committee (Technische Universität Berlin, Germany) and all participants gave written informed consent in accordance with the Declaration of Helsinki.

#### 4.1.2 Experimental Paradigm

Participants were situated in a virtual environment that displayed only floor texture. They were instructed to follow a sphere that rotated around them and stopped unpredictably on a trial at different eccentricities. The task of participants was then to rotate back to indicate their initial heading direction. The task was self-paced and participants initiated a trial with a button press with their index finger. Each trial started with the appearance of a red pole indicating the starting position participants had to face. After signaling alignment with a second button press, the pole disappeared and a red sphere appeared, circling around the participant in a distance of 30m. Participants rotated on the spot to keep the sphere in the center of their view. The sphere stopped and turned blue to mark the end of the outward rotation. Participants then rotated back and indicated their estimated initial heading by a button press. Participants rotated both clockwise and counter-clockwise, in varying velocities and eccentricities (30° to 150°), in a randomized order, summing up to 140 trials. The task was completed twice, once using a traditional 2D monitor setup where movement was controlled through a joystick (stationary condition), and once with a virtual reality setup where movement was controlled through physical body movement (mobile condition). The order was balanced across participants. An overview of the paradigm can be seen in figure 1.

**FIGURE 1.**
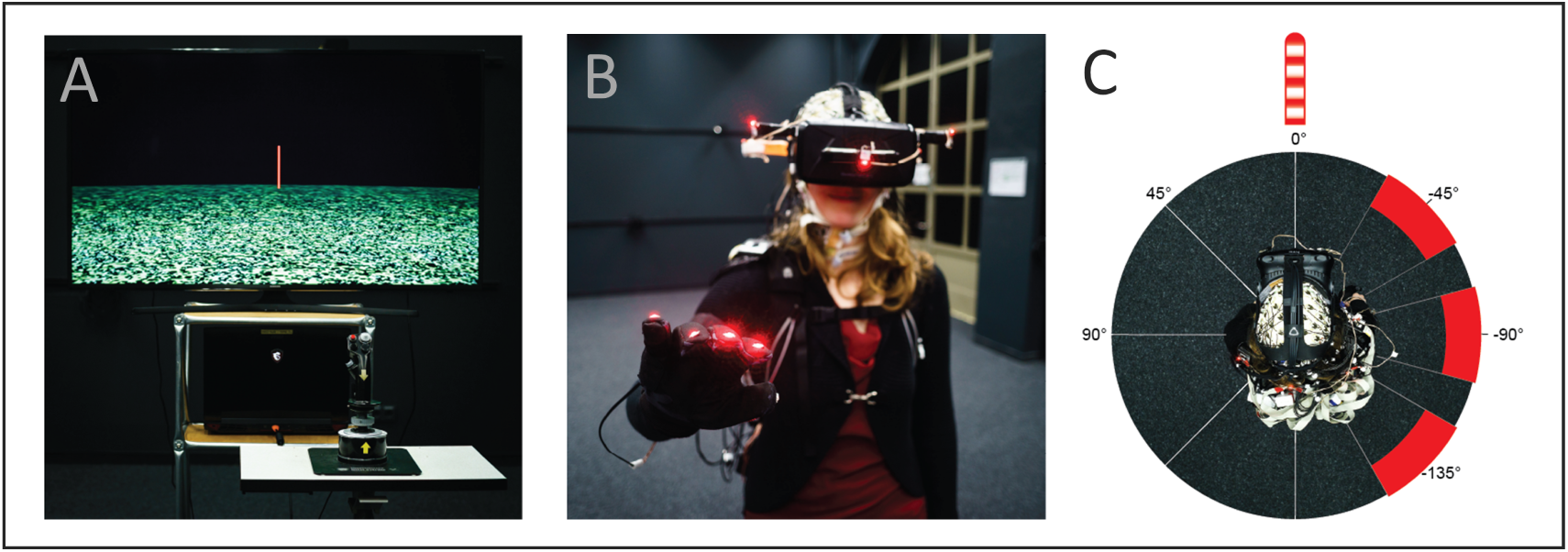
Experimental setup and paradigm. **A** Setup of the stationary condition with joystick rotation (visual flow only), displaying a sparse virtual environment with a local landmark providing the initial heading direction (pole). The joystick was placed on a table in front of the standing participant. **B** A Mobile Brain/Imaging setup with a participant wearing a head-mounted virtual reality (VR) display, high-density EEG including an EMG neckband, and motion capture devices (red LEDs on gloves and VR). **C** Top-down view of a participant in the mobile condition, displaying the rotation eccentricities (varying +/- 15° around 45°, 90°, and 135°, respectively).

In the stationary condition, participants stood in front of a TV monitor (1.5m distance, 40” diagonal size, HD resolution, 60 Hz refresh rate) and were instructed to move as little as possible. In the mobile condition, they were wearing a head-mounted virtual reality display (HTC Vive, 110° field of view, 2×1080×1200px resolution, 90 Hz refresh rate) and a backpack PC so no cables constrained their movement, and completed the task by physically rotating on the spot. Each condition was preceded by a baseline of three minutes during which participants were asked to stand still, keep their eyes open, and to look straight ahead. Completing each condition took around 30 minutes, where the mobile condition was slightly shorter than the stationary condition due to faster physical rotations during the response movement.

#### 4.1.3 Data Recording

In both conditions, EEG was recorded from 157 active electrodes on both the scalp (129 electrodes) and neck of the participant (28 electrodes). The latter were used to specifically record neck muscle activity for a potential benefit in data cleaning. Electrodes on the scalp were placed using an elastic cap with an equidistant design. The electrodes on the neck were placed with a custom design neckband (EASYCAP, Herrsching, Germany). All channels were referenced to an electrode close to the standard FCz position and data was recorded with a sampling rate of 1000 Hz. The data was band-pass filtered from 0.016-500 Hz (BrainAmp Move System, Brain Products, Gilching, Germany) and impedances were kept below 10kΩ for electrodes on the scalp, and below 50kΩ for neck electrodes. Individual electrode locations were recorded using an optical tracking system (Polaris Vicra, NDI, Waterloo, ON, Canada).

In addition to the EEG, motion capture data was recorded using either the camera location in the virtual environment, or, in the mobile condition, the VR lighthouse tracking system (HTC Corporation, Taoyuan, Taiwan) of the head-mounted display, and active LEDs on the feet, around the hip, and on the shoulders with the Impulse X2 System (PhaseSpace Inc., San Leandro, CA, USA), all with a sampling rate of 90 Hz. Motion capture data was not used for the analyses presented here. Data and event marker streams of different sources were time-stamped and recorded using Lab Streaming Layer ^1^.

### 4.2 Processing Pipeline

The data was analyzed in MATLAB (R2016b version 9.1; The MathWorks Inc., Natick, Massachusetts, USA), using custom scripts based on the EEGLAB toolbox (Delorme and Makeig, 2004, version 14.1.0). We investigated the effects of different factors on the quality of the resulting ICA decomposition. To this end, we systematically assessed the impact of the experimental protocol (stationary vs. mobile condition), the channel density (5 different montages), and the high-pass filter cutoff frequency (from no filter up to 4 Hz cutoff). A schematic overview of the data processing pipeline can be seen in figure 2.

**FIGURE 2.**
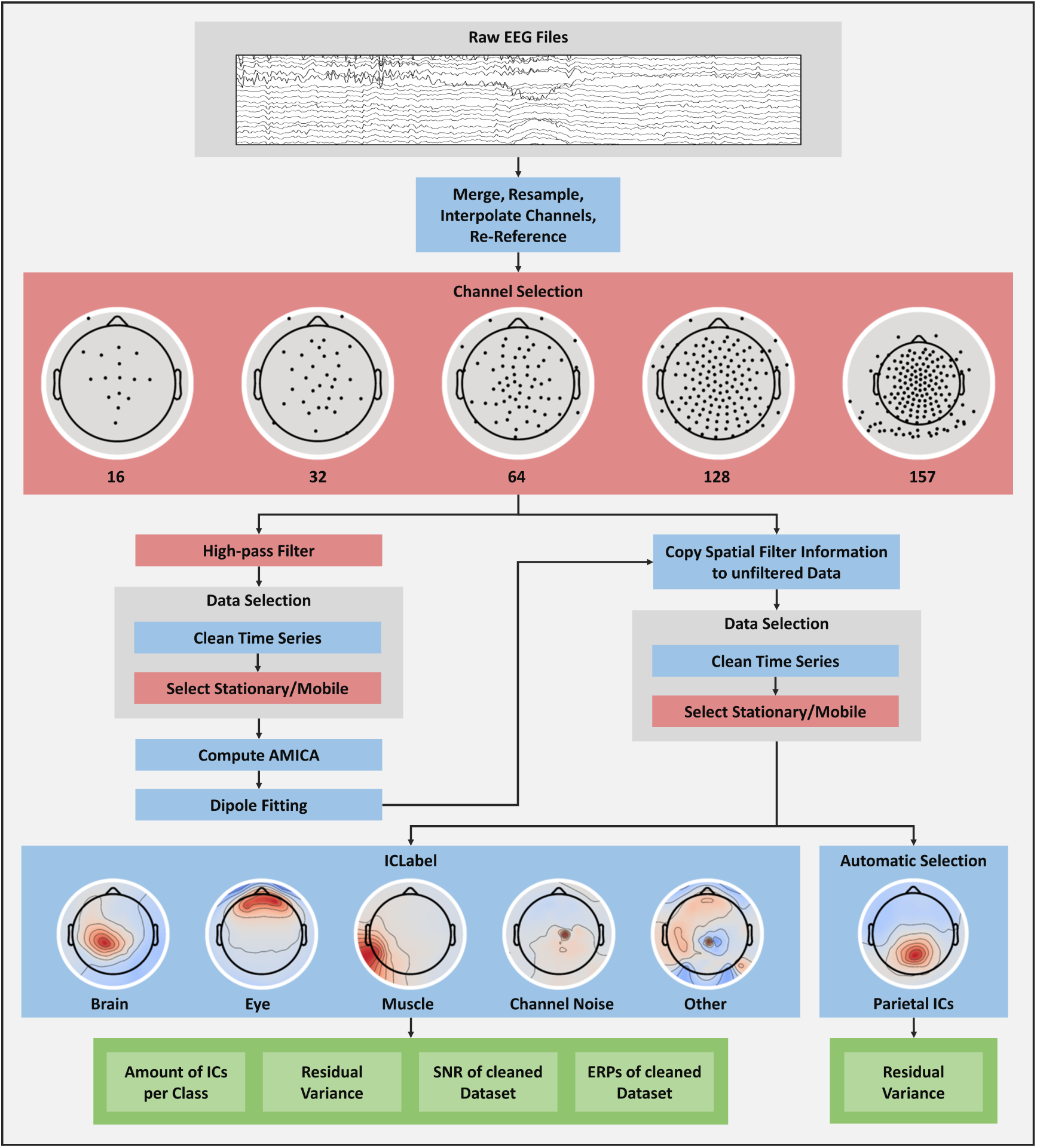
Schematic overview of the processing pipeline. Blue boxes mark processing steps which were executed identically for all datasets, red boxes mark a selection of conditions, green boxes mark final quality measures. Steps are described in section 4.2 - *Processing Pipeline*.

#### 4.2.1 Preprocessing

Data from both conditions was first appended and individual channel locations were loaded. Raw EEG data was then lowpass filtered at 125 Hz, resampled to 250 Hz and manually scored for bad channels, which were subsequently interpolated. Lastly, channels were re-reference to the average reference.

#### 4.2.2 Channel Selection

In the next step, we selected channels of the dataset to be included in the analyses. This were either all channels, using the full equidistant setup with 129 scalp and 28 neck electrodes or a roughly equidistant subset of only the scalp electrodes, resulting in a 128, a 64, a 32, and a 16 channel scalp setup. Since the data contained free eye movements, the two electrooculogram (EOG) electrodes below the eyes were kept for all setups except the 16 channels, where one of them was dropped. See figure 2 for a visualization of the channel layout. These data constituted the basic datasets (“datasets A”).

#### 4.2.3 High-pass Filtering

In order to compare the impact of different high-pass filter frequencies on ICA decompositions, the five datasets A were filtered with a zero-phase Hamming window FIR-filter (EEGLAB firfilt plugin, version 1.6.2) with varying cutoff frequencies. In many cases, it is advisable to specify the filter order in detail to achieve maximal control of the process (see Widmann et al. (2015) for a practical guide to filtering EEG data). The filter passband-edge defines where signal attenuation begins, the cutoff frequency is the frequency where the signal is attenuated by 6 db and can be regarded as the frequency where the filter starts to have a noticeable effect. The transition bandwidth is double the difference between passband-edge and cutoff frequency and is specified by the filter order. The stopband-edge is the passband-edge minus the transition bandwidth and can be regarded as the frequency where the signal attenuation reaches its full effect. At this point, it should be noted that in EEGLAB filters are specified by passband edge and follow a heuristic to find a suitable filter order (and thus transition bandwidth) depending on the frequency. For example, a default 1 Hz filter as used by EEGLAB routines has a transition bandwidth of 1 Hz and a cutoff of 0.5 Hz, whereas a 3Hz filter has a transition bandwidth of 2 Hz and a cutoff frequency of 2 Hz. For the present study, we used a constant filter order of 1650 to ensure comparability, resulting in a transition bandwidth of 0.5 Hz independently of the passband-edge^2^. This means that a filter with a specified passband edge of 1 Hz and a transition bandwidth of 0.5 Hz leads to a cutoff frequency of 0.75 Hz and a stopband frequency of 0.5 Hz. In the further course of this paper, we use the cutoff frequency to specify the filter. As the literature suggests, we focused our analysis on lower frequencies. Since the transition bandwidth was 0.5 Hz, the lowest cutoff frequency that could be applied was 0.25 Hz. We then increased the frequency in steps of 0.25 Hz for lower frequencies up to 1.5 Hz, then in steps of 0.5 Hz up to 3 Hz, and added a 4 Hz filter as the highest frequency. Additionally, we added an analysis without any filtering (0 Hz) for comparison. This resulted in 11 different filter settings for all of the datasets A.

#### 4.2.4 Data Selection

After filtering, a manual cleaning followed where the data was scored for strong artifacts. The marked timepoints were saved and rejected from all filtered datasets. The selection of the stationary and mobile experimental conditions was made based on the event markers present in the data. Importantly, to ensure comparability, both the stationary and mobile conditions had to be of the same length. As a consequence, the longer dataset was cut to the length of the shorter dataset (on average, datasets were 27 min long, SD = 5.8 min). Overall, this resulted in 110 datasets per subject composed of 5 (channel montages) x 11 (filter cutoff) x 2 (movement condition) that entered an ICA decomposition.

#### 4.2.5 Independent Component Analysis

All final 2090 datasets were then decomposed using the AMICA algorithm (Palmer et al., 2011). AMICA was chosen since it is considered the best ICA algorithm (see section 2.1 - *Achieving an Optimal Decomposition*) and is widely used by different research groups. Although the impact of filtering has been evaluated for algorithms other than AMICA, AMICA itself was not often subject to these investigations. We used one model and ran AMICA for 2000 iterations on all datasets. Since we interpolated channels previously and used an average reference for our datasets, we also let the algorithm perform a principal component analysis rank reduction to the number of channels minus 1 (average reference) minus the number of interpolated channels. All computations were performed using four threads on machines with identical hardware, an AMD Ryzen 1700 CPU and 32GB of DDR4 RAM. Overall, computation time amounted to 4340 hours for all participants and datasets.

#### 4.2.6 Dipole Fitting

Subsequently, for every resulting IC, an equivalent dipole model was computed as implemented by the DIPFIT plugin for EEGLAB. For this purpose, the individually measured electrode locations of every participant were warped (rotated and rescaled) to fit a boundary element head model based on the MNI brain (Montreal Neurological Institute, MNI, Montreal, QC, Canada). The dipole model includes an estimate of IC topography variance which is not explained by the model (residual variance, RV).

#### 4.2.7 Transfer of AMICA and Equivalent Dipole Model Structures

Since the final quality measures of the resultant AMICA decomposition needed to be computed on comparable unfiltered data to allow for a direct comparison of ICs, we copied the results of the AMICA and equivalent dipole model fitting back to dataset A. Especially the automatic IC classification based on ICLabel (Pion-Tonachini et al., 2019) proved to be sensitive to filtering in the lower frequencies in prior tests. Subsequently the data was then cleaned and split into the conditions again (identical to section 4.2.4 - *Data Selection*).

#### 4.2.8 Automatic Component Classification

The next part of the processing was the automatic classification of ICs using the ICLabel algorithm (Pion-Tonachini et al., 2019). ICLabel is a classifier trained on a large database of expert labelings of ICs, which classifies whether or not ICs are of brain or non-brain origin, including eye, muscle, and heart sources as well as channel and line noise artifacts and a category of other, unclear, sources. The class probability is provided allowing both for a more fine-grained analysis of probabilities and a classic popularity vote classification. Classifying based on a class probability threshold per class can be beneficial when the focus of interest lies mainly on one class, but it can also lead to ICs which have zero or more than one class labels assigned. Since we were interested in comparing the different classes, we used the popularity vote for our analysis. As a result, ICs received the class with the maximal probability as their class label. Two versions of the ICLabel algorithm exist: i) the *default* version which uses the IC activity spectrum (1-100Hz), IC topography, and IC activity autocorrelation as features for classification, and ii) the *lite* version which does not take autocorrelation into account. The latter is faster to compute and uses less RAM, especially for larger datasets, and although the classification of brain ICs can be slightly better in the default version, classification of other sources like eyes and muscles can be better using the lite version. Hence, we ran ICLabel twice using both versions but focus our analysis on the lite version^3^. See figure 2 for example patterns of the most important classes.

#### 4.2.9 Automatic Selection of Parietal Components

In addition to the ICLabel classification, we automatically selected one parietal IC for each decomposition, based on a topographic weight map. To allow automatic selection of this parietal component, we took the first 10 + number of ICs / 3 ICs with a RV of <0.1 into account. The additional inspection of a specific brain IC allowed for the investigation of the impact of the preprocessing independently of ICLabel. Additionally, this allowed for investigating the effect of channel density, filtering and movement on specific scalp topographies as opposed to a general decomposition quality. This can be important when using ICA to examine the data on the source level, for example in a parietal region of interest. See figure 2 for an example parietal pattern.

### 4.3 Quality Measures

In order to compare the decomposition quality, we extracted several features addressing both general and practical considerations. First, we considered the ICLabel classifications. The focus of most EEG research lies on brain signal analysis and the removal of other sources that are considered artifactual contributions. In MoBI research, in contrast, analyzing muscle and eye activity as signals can be very important to make sense of the data and potentially to be used as a source of insight into cognitive processes. Hence, we were interested in the amount of brain, eye, and muscle ICs as signal sources, and the amount of other Ics as a proxy of general decomposition quality.

Additionally, we were interested in the residual variance of ICs after fitting an equivalent dipole model. The RV, especially of brain components, is an important measure to estimate the quality of a component (Delorme et al., 2012). A low RV means that the respective independent component is largely dipolar in nature, which in turn means it is more physiologically plausible and more likely to be of brain origin, since the standard head models only include dipoles in the cortex. Often, this measure is used to separate brain ICs from other ICs where ICs with an RV <0.15 are treated as more likely originating in the brain. We were interested in the mean intra-class RV for the ICLabel classes as well as the mean RV of the parietal ICs.

Finally, as a practical measure for researchers, event-related-responses (ERPs) were computed for all datasets and further examined on their the signal-to-noise ratio (SNR). To this end, the data was pruned with ICA by removing all ICs that were classified as non-brain classes and only brain ICs were backprojected to the sensor level. Since the data was previously scored for artifacts in the time domain, only trials which were comparably artifact-free entered the ERP. The ERPs were computed at an electrode in our equidistant layout which was positioned closest to the POz electrode in the standard layout (POz’). Importantly, to not distort the results we used no frequency filter on the data (as the ICA results were copied back to the unfiltered data), but only the spatial filter of the ICA. We then extracted epochs (−600 ms to +1200 ms) around the trial-onset event (onset of the moving sphere) for which we expected a parietal P300 to occur, and removed the pre-stimulus baseline activity. The two mobility conditions (stationary, mobile) did not contain the same amount of events, as the stationary condition had to be cut short to fit in length to the mobile condition in which participants rotated back faster and thus were able to answer more trials within the same time. To ensure comparability between the conditions, we determined the minimal number of available events for both movement conditions per subject and used this number of events in both conditions to compute the ERPs. Epochs were averaged per subject and condition (channel density, high-pass filter before computing ICA, movement condition), and the final measures for signal and noise were computed. The mean amplitudes from 250 ms to 450 ms served as the signal which was divided by the standard deviation in the 500 ms pre-stimulus interval to compute the SNR (Debener et al., 2012).

## 5 RESULTS

Overall, the ICA decompositions were sensitive to the different preprocessing parameters. Figure 3 shows the results for the number of ICs in the *Brain, Muscle, Eye*, and *Other*, classes, as well as their mean RV. The results of the RV values of the parietal ICs can be seen in figure 4. Finally, the practical quality measures of ERPs and SNR values can be found in figure 5.

**FIGURE 3.**
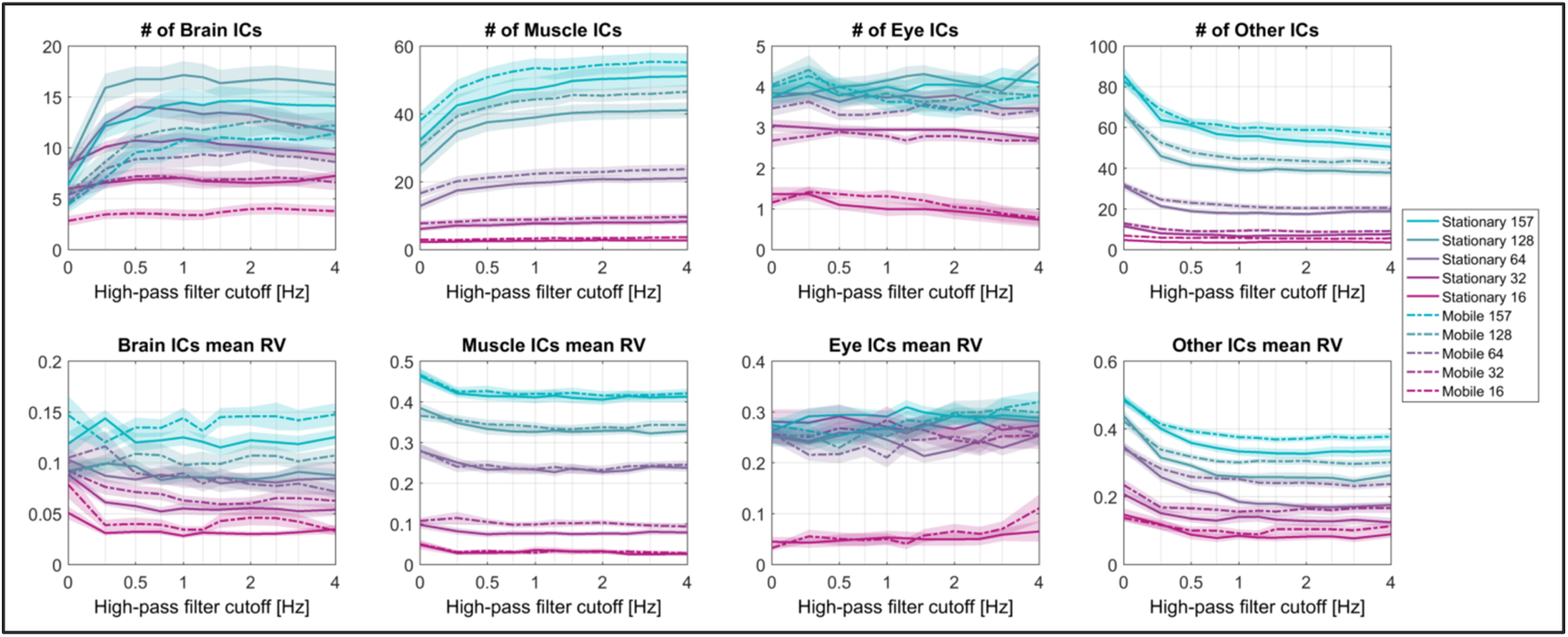
Results for the ICLabel classifications. Top row: amount of ICs per class, bottom row: mean residual variance (RV) per class. 0 Hz refers to no filter being applied before computing ICA. Note the logarithmic scaling of the abscissa with grid lines for each available filter frequency.

**FIGURE 4.**
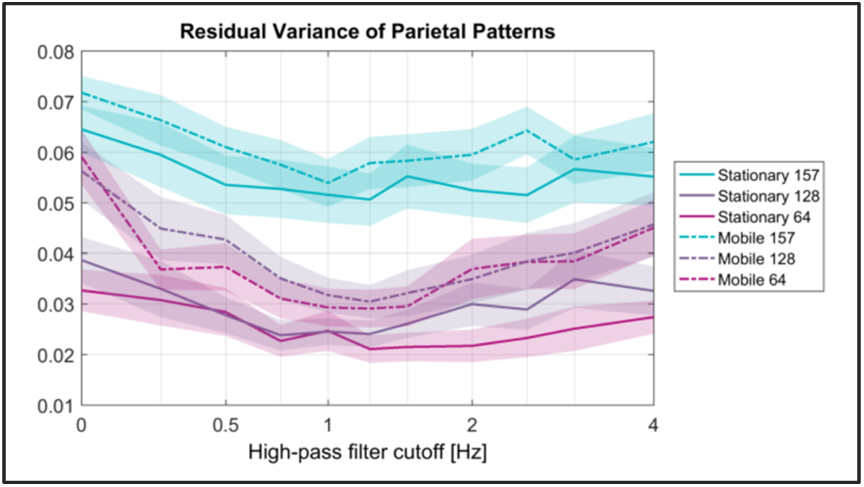
Residual variance (RV) of the parietal patterns. Only channel montages of 64 and more channels were considered. 0 Hz refers to no filter being applied before computing ICA. Note the logarithmic scaling of the abscissa with grid lines for each available filter frequency.

**FIGURE 5.**
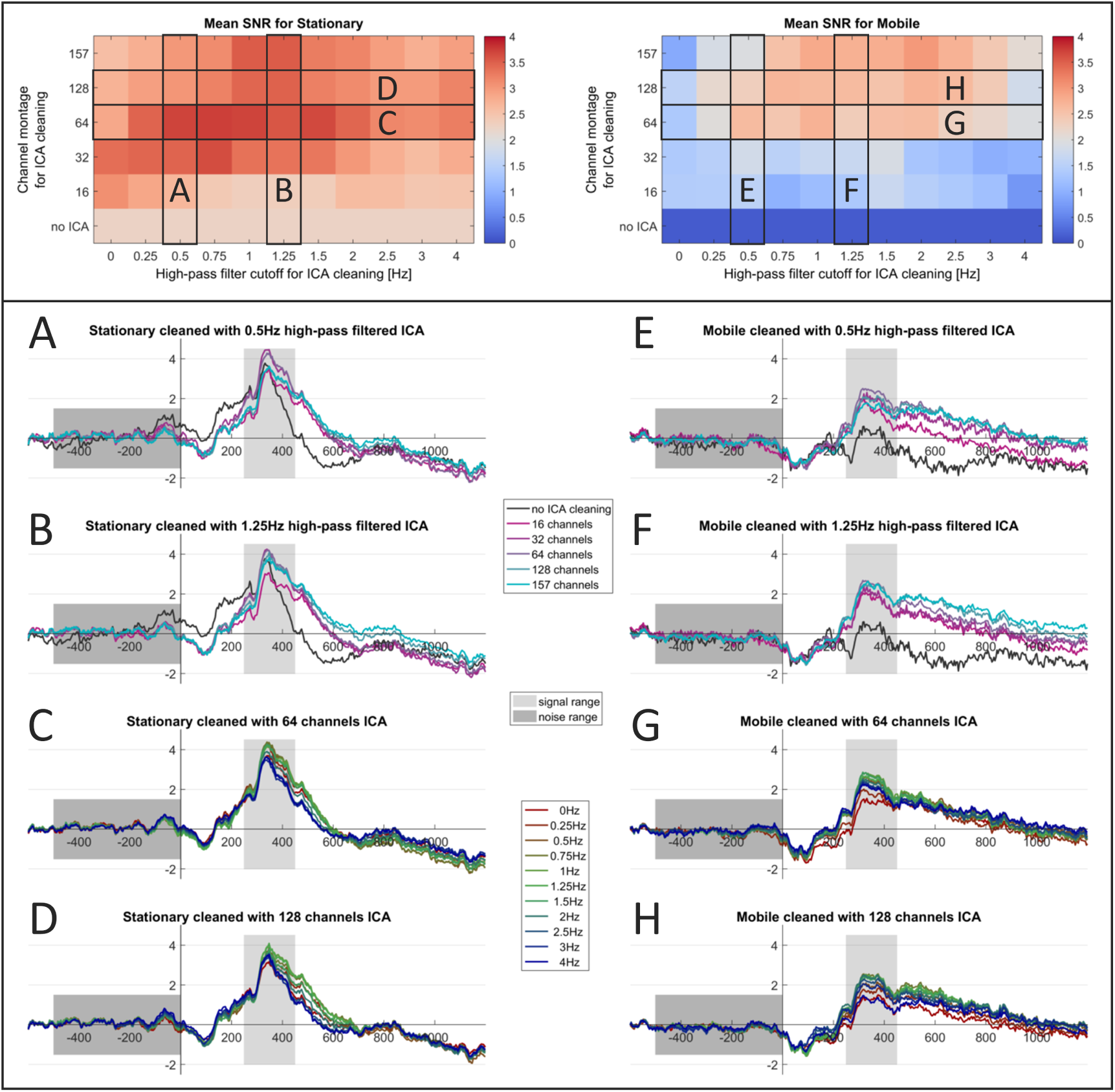
Practical quality measures. **Top:** Signal-to-noise ratios (SNRs) of the event-related-responses (ERPs) computed on uncleaned data and data that was cleaned with ICA by removing all non-brain ICs as classified by ICLabel. ERPs were computed on the POz’ electrode for the trial-onset event. Note that as the ICA results were copied back to the unfiltered datasets prior to ERP computation, the datasets themselves only differed in the ICA decompositions that were used for cleaning, no frequency filter was applied before computing the ERPs. SNR was defined as the mean amplitude in the 250 ms - 450 ms interval divided by the standard deviation in the 500 ms pre-stimulus interval. 0 Hz refers to no filter being applied before computing ICA. **Bottom:** Corresponding ERPs, plotted either for different channel montages and a fixed filter cutoff frequency used before computing ICA (columns of SNR plots, A, B, E, F), or different cutoff frequencies and a fixed channel montage (rows of SNR plots, C, D, G, H).

Clear differences could be observed between the stationary and mobile data. The stationary data contained more brain ICs and less muscle ICs than the mobile data, additionally, the mobile data contained more ICs classified as “other”. Interestingly, the number of eye ICs did not differ between the mobility conditions. A larger RV of brain ICs could be observed in the mobile condition, however, this difference was not very large (around 0.03) and especially for the montages with 128 channels it was small or non-existent. Considering the parietal ICs specifically, however, the mobile condition consistently exhibited a higher RV value than the counterparts in the stationary condition (around 0.01 difference). The SNR of the ERPs was considerably greater in the stationary condition than in the mobile condition, and the shape of the ERP in the two mobility conditions was different, where a larger P300 peak with a steeper offset occurred in the stationary condition.

The ICA decomposition was also clearly influenced by the channel montage, with generally more ICs being present in each class in higher channel densities. This was evident also in the number of brain ICs, but even with 16 channels there were still ICs classified as brain. Importantly, the difference in brain ICs between the channel montages was less pronounced than the difference in muscle and other ICs, thus showing a more pronounced stability of the brain ICs. Note that the maximum of brain ICs was not reached with the montage containing the neck band (157 channels), but with 128 scalp channels. This was true for both mobility conditions, but the detrimental effect of the neck band was less strong in the mobile condition. The number of eye ICs was stable across channel montages, with densities of 64 channels and upward containing four eye ICs, the 32 channel montage containing three eye ICs, and only the 16 channel montage containing only one IC classified as reflecting eye movement activity. The RV of channel montages with fewer channels was lower in general. This is evident in all classes except the eye ICs, where only the 16 channel montages showed a decrease in RV. The 16 channel montages reached RV values of <0.1 in most cases, including not only physiological, but also other ICs. The difference between the channel montages were more pronounced for muscle and other ICs than for brain ICs, which means that the difference in RV values between brain and non-brain ICs was larger when using more channels. In the parietal ICs, the 157 channel montage showed a general increase of RV (around 0.025) above the 64 and 128 channel montages, whereas only a slight increase could be observed in the 128 channel montage over the 64 channel montage. Interestingly, when looking at the SNR values and the ERP waveforms, the 64 channel montage already lead to very good or the best results and cleaning the data with 16 channels already led to a substantial improvement in SNR. Visual inspection of the ERPs also showed clear improvements already with 16 channels, especially in the mobile condition where the uncleaned data contained almost no signal.

The high-pass filter applied before computing ICA had a considerable influence as well. For brain, muscle, ICs, an increase of the number of ICs in these classes with increasing high-pass frequencies could be observed, especially when compared to no filter (0 Hz). The number of eye ICs appeared to be insensitive to high-pass filtering, whereas the number of other ICs dropped with increasing filter frequency. The effect of filtering also exhibited a ceiling or floor effect, where a filter higher than 1-2 Hz mostly did not affect the results any further. Muscle ICs increased in numbers a little slower, approaching an optimum from 2Hz onward and continued to increase even up to 4 Hz cutoff (maximum filter applied). The RV of brain ICs appeared relatively stable across different filter frequencies, implying that an increased number of brain ICs did not coincide with classifying less dipolar ICs as originating in the brain. The mean RV of brain ICs ranged from 0.03 (stationary condition with 16 channels) to 0.15 (mobile condition with 157 channels). RV values of muscle and other ICs droppped with increasing filter frequency up to 1 Hz, which was more noticeable with higher channel densities. In the parietal ICs, a slightly different pattern could be observed, where the RV did not approach a floor asymptote, but increased again after reaching the minimum around 1 Hz. This inverted-U shape with increasing filter frequencies could also be observed in the SNR measures of the ERPs and the ERP waveforms themselves, where mid-range filters show a larger P300 signal, not only in the range used for the SNR (250 ms to 450 ms post-stimulus) but continuing until around 600 ms post-stimulus.

Importantly, the effect the filter frequencies had on the ICA decomposition was less pronounced with decreasing number of channels, with the 16 channel montage exhibiting almost no change between filter frequencies any more. In the parietal ICs, the already small difference between the 64 and the 128 channel montage disappeared at the optimal filter frequency. The SNR in higher channel densities generally required a higher filter to reach its maximum which could also be seen in the ERPs themselves.

Finally, the effects of filter and mobility conditions appeared to interact as well. For the number of brain ICs, the necessary filter to reach the maximum showed a marked difference between stationary and mobile data: The maximum number of 17.2 brain ICs was reached in the stationary condition with 128 channels and a 1 Hz filter, the mobile condition reached its maximum of 12.7 brain ICs also with 128 channels, but a 2.25 Hz filter. This pattern could also be found in the RV of parietal ICs, where the RV reached its minimum earlier in the stationary condition (0.024 with 0.75 Hz and 128 channels) than in the mobile condition (0.030 with 1.25 Hz and 128 channels). The SNR of the ERPs also showed that in the mobile condition a higher filter was necessary to obtain the best results. As expected from the SNR values, the ERP waveforms with the highest P300 values in the mobile condition were the ones cleaned with ICA computed on higher filtered data, especially noticeable with 128 channels where the maximal P300 occurred with the 2.25 Hz filter in the mobile condition as opposed to 1.25 Hz in the stationary condition.

## 6 DISCUSSION

EEG is a widely adopted tool in neuroscientific research and in the recent years new trends towards more active and mobile experiments emerged, allowing for the investigation of more natural cognitive processes “in the wild”. These experiments, however, come with the drawback of additional and stronger artifactual contributions in the data which can mask electrical activity originating from the brain. Separating the different sources is thus a key step in modern EEG research and does not only allow for an analysis of clean data but also an estimation of the source activity and their cortical origins. Although blind source separation techniques like ICA are widely adopted as a tool to achieve this goal, the influence of different factors on this decomposition is not always clear. In this study we investigated the impact of movement of participants, channel density, and high-pass filter cutoff frequency during preprocessing on the decomposition of EEG data with ICA. We evaluated the outcome of ICA based on differently preprocessed data using the number of brain, muscle, eye, and other ICs as classified by ICLabel, their dipolarity, and the SNR of ERPs on the cleaned data.

The results show that, as expected, participant movement has a detrimental effect on the decomposition and generally leads to fewer and less dipolar brain ICs but more muscle ICs in the data. This is not surprising as more artifacts and especially more muscle activity is present in MoBI data which take up degrees of freedom for the ICA decomposition. Importantly, ICA is still a powerful tool for cleaning EEG data even in light of increasingly noisy recordings from MoBI and mobile EEG experiments, as the effect of cleaning on SNR and ERPs shows. In fact, in mobile EEG studies it is possible to miss the entire signal of an ERP without cleaning the data, but it may be reconstructed by removing non-brain ICs. Analyzing MoBI data with ICA or a comparably powerful cleaning method thus appears to be vital. It is to note that the strong difference of the ERP waveform between stationary and mobile data is not necessarily an effect of artifacts and increased noise alone. Since the brain needs to fulfill a variety of additional tasks when moving the body by preparing and executing motor commands with constant sensory feedback, attention to a specific stimulus may be limited (Ladouce et al., 2019) and decreasing sensory mismatch might change oscillatory processes (Gramann et al., 2018).

Considering the effect of the number of EEG channels, it can be stated that a higher scalp electrode density generally leads to a better ICA decomposition. However, there seems to be a ceiling effect when cleaning the sensor data with ICA which is reached already when using 64 channels, as shown in the ERP SNR analysis of both mobility conditions. To our surprise, even an ICA with 16 channels is powerful enough to reconstruct the ERP to a useful degree and is a noticeable improvement over not using ICA at all, especially in the mobile case. Of course, this does not mean that a higher-density data recording does not improve the results of a decomposition. Using more channels results in more brain ICs, which in turn will lead to the possibility of a more detailed source-level analysis, making EEG a powerful tool to truly image the brain in action. A second surprise was the observed detrimental effect of an EMG neckband on the number of brain ICs and their RV. This might have two reasons: First, it could be that the neckband is not an ideal candidate for measuring EMG activity. As 28 electrodes are placed around the neck, the width of the band might have been too large in some subjects, leading to movement of the neck band and the included electrodes and thus changes in the electrode-skin contact. Additionally, since the neckband was fixed, turning the head might have led to electrodes shifting over the skin, leading to artifacts and EMG measurements that were potentially spatially unstable, introducing non-stationarity into the ICA decomposition and thus violating assumptions of the ICA model. In sum, the EMG neckband that was used might have introduced more artifacts to the data than adding useful information and degrees of freedom for the spatial filter. However, another explanation could be that since ICLabel uses images of the IC topography as a feature for classification, the change in the topography due to the added neck channels (see figure 2) led to less accurate classifications and thus more missed brain ICs. When looking at the SNRs and ERPs, although not being particularly helpful, the neckband seemed to be unproblematic, and especially in the mobile case the highest SNR was reached with the 157 channel montage. In light of these considerations and the beneficial effect of EMG found in simulation studies (Richer et al., 2019), it needs to be further examined whether using sticky electrodes for recording neck muscle activity instead of a neckband improves the results. One other observation we made is that the RV of ICs decreases with fewer channels, seemingly suggesting a better, more dipolar, decomposition. This effect could be caused by an actually better decomposition due to more samples available relative to the number of channels (as the dataset size was kept identical to ensure that the same information entered the ICA). On the other hand, it might be caused by less measurement points (channels) available to compute the RV in these recordings. Extrapolating to the case of a single-channel recording, no RV would be measurable any more. Exploring this factor by adjusting the dataset length to the number of channels is an important option for future investigations. Independently of the underlying cause, however, it is important to note that in experiments of typical lengths of 30 to 60 minutes, RV may be only useful to dissociate brain and non-brain ICs when recording with higher-density montages of 64 channels and more.

Lastly, we were able to confirm our hypothesis that high-pass filtering before computing ICA does improve the decomposition and a filter cutoff of 0.5-2 Hz is optimal. However, we found no single optimal filter, instead, the filtering should be adjusted depending on other factors of the experiment. In standard stationary experiments with 64 channels a high-pass filter cutoff of 0.5 Hz is acceptable, but with increasing number of channels a higher filter cutoff of up to 1.25 Hz is necessary to achieve the best decomposition. This effect was even more pronounced in the mobile condition where the decomposition improved further with cutoff frequencies of up to 2 Hz. Interestingly, even though we come to similar conclusions as Winkler et al. (2015) regarding the filter cutoff, we did observe clear changes in the ERP waveforms which the authors did not find. We believe this could be due to the experimental paradigm, as even in our stationary condition participants were standing upright controlling the visual flow with a joystick. Thus there were likely more artifacts present than in a classic auditory oddball paradigm with seated participants and the effect of cleaning the data with ICA was more noticeable. Comparing our results to those of Frølich and Dowding (2018), we could not confirm that a very high cutoff frequency led to better results, as we saw detrimental effects after reaching the optimal filter cutoff. These conflicting results may be due to the fact, that Frølich and Dowding (2018) employed a 45 Hz low-pass filter even though they specifically investigated muscle activity, which is more prevalent in higher frequencies. We want to emphasize again that when discussing filters in this paper we used the cutoff frequency, not the passband-edge as the defining parameter, and when using EEGLAB it is recommended to specify the correct filter (see section 4.2.3 - *High-pass Filtering*).

We conclude that obtaining an optimal ICA decomposition when analyzing EEG data is highly relevant, not only for source-level analysis but also for cleaning sensor data, and it is especially effective and necessary when expecting more and stronger artifacts. We would like to finalize this paper by providing some recommendations as a set of “best practices” when performing ICA on EEG data. First of all, when computing ICA to remove eye and muscle artifacts it is important to do this on data which was high-pass filtered but not low-pass filtered, and and it is unproblematic to apply the obtained decomposition to unfiltered data for further analysis. Second, higher-density recordings of 64 and more channels should be used when aiming for an optimal recovery of the brain signals and especially when doing source-level analysis, as low-density recordings can not separate neural sources adequately. Third, an increasing channel density is required with increasing movement range and velocity in the experimental protocol. Fourth, when no high-density recording is possible, ICA can still be used to clean the sensor data from eye and muscle activity artifacts. Last, but not least, we recommend using higher high-pass filter cutoffs than traditionally used. While 0.5 Hz might be acceptable for 64 channels in stationary experiments, using a 1 Hz filter is not detrimental and ensures a good decomposition also for higher-density recordings with more noise being present in the data. For MoBI experiments with significant noise even higher filters of 1.5 or even 2 Hz should be employed before computing ICA, depending on the channel montage.

## ACKNOWLEDGEMENTS

This work was supported by the DFG (DFG-Project GR2627/8-1) and USAF (ONR 10024807). We wish to thank Jonna Jürs and Yiru Chen who assisted in collecting the data, and Emma Auerbach Brode for her initial contributions to the project. We also sincerely thank Olaf Dimigen for important notes on filter specifications.

## CONFLICT OF INTEREST

The authors declare no conflict of interest.

## AUTHOR CONTRIBUTIONS

M.K. and K.G. designed the research; M.K. participated in the original data collection; M.K. performed the data analysis and wrote the first draft of the paper; K.G. and M.K. edited the paper.

## DATA ACCESSIBILITY

Data relating to these experiments are available upon request from the corresponding author.

## ABBREVIATIONS

EEG: electroencephalography;
EOG: electrooculography;
ERP: event-related-response;
EXG: electrooculography and electrocardiography;
IC: independent component;
ICA: independent component analysis;
MEG: magnetoencephalography;
MoBI: Mobile Brain/Body Imaging;
RV: residual variance;
SNR: signal-to-noise ratio;
VR: virtual reality;

## Footnotes

1 https://github.com/sccn/labstreaminglayer

2 This can be reproduced in MATLAB/EEGLAB with `[EEG, com, ∼] = pop_eegfiltnew(EEG, highpassPassbandEdge, 0, 1650, 0, [], 1`), note that in EEGLAB the specified value is the passband edge, not the cutoff frequency, which in this case is `desiredCutoff + 0.25`.

3 A comparison of the two algorithms can be found in the supplementary material.

## Notes

Funding information This work was supported by the DFG, (DFG-Project GR2627/8-1) and USAF (ONR 10024807).

### Competing Interest Statement

The authors have declared no competing interest.

## REFERENCES

Artoni, F., Menicucci, D., Delorme, A., Makeig, S. and Micera, S. (2014) RELICA: A method for estimating the reliability of independent components. NeuroImage, 103, 391–400.

Bell, A. J. and Sejnowski, T. J. (1995) An information-maximization approach to blind separation and blind deconvolution. Neural computation, 7, 1129–1159.

De Sanctis, P., Butler, J. S., Malcolm, B. R. and Foxe, J. J. (2014) Recalibration of inhibitory control systems during walking-related dual-task interference: A Mobile Brain-Body Imaging (MOBI) Study. NeuroImage, 94, 55–64.

Debener, S., Minow, F., Emkes, R., Gandras, K. and de Vos, M. (2012) How about taking a low-cost, small, and wireless EEG for a walk? Psychophysiology, 49, 1617–1621.

Delorme, A. and Makeig, S. (2004) EEGLAB: An open source toolbox for analysis of single-trial EEG dynamics including independent component analysis. Journal of Neuroscience Methods, 134, 9–21.

Delorme, A., Palmer, J., Onton, J., Oostenveld, R. and Makeig, S. (2012) Independent EEG sources are dipolar. PLoS ONE, 7, e30135.

Dimigen, O. (2020) Optimizing the ICA-based removal of ocular EEG artifacts from free viewing experiments. NeuroImage, 207, 116117.

Djebbara, Z., Fich, L. B., Petrini, L. and Gramann, K. (2019) Sensory-motor brain dynamics reflect architectural affordances. Proceedings of the National Academy of Sciences, 1–31.

Ehinger, B. V., Fischer, P., Gert, A. L., Kaufhold, L., Weber, F., Pipa, G. and König, P. (2014) Kinesthetic and vestibular information modulate alpha activity during spatial navigation: a mobile EEG study. Frontiers in human neuroscience, 8, 71.

Frølich, L. and Dowding, I. (2018) Removal of muscular artifacts in EEG signals: a comparison of linear decomposition methods. Brain Informatics, 5, 13–22.

Gehrke, L., Iversen, J. R., Makeig, S. and Gramann, K. (2018) The invisible maze task (imt): Interactive exploration of sparse virtual environments to investigate action-driven formation of spatial representations. In Spatial Cognition XI (eds. S. Creem-Regehr, J. Schöning and A. Klippel), 293–310. Cham: Springer International Publishing.

Gramann, K., Ferris, D. P., Gwin, J. and Makeig, S. (2014) Imaging natural cognition in action. International Journal of Psychophysiology, 91, 22–29.

Gramann, K., Gwin, J. T., Bigdely-Shamlo, N., Ferris, D. P. and Makeig, S. (2010) Visual evoked responses during standing and walking. Frontiers in Human Neuroscience, 4, 202.

Gramann, K., Gwin, J. T., Ferris, D. P., Oie, K., Jung, T. P., Lin, C. T., Liao, L. D. and Makeig, S. (2011) Cognition in action: Imaging brain/body dynamics in mobile humans. Reviews in the Neurosciences, 22, 593–608.

Gramann, K., Hohlefeld, F. U., Gehrke, L. and Klug, M. (2018) Heading computation in the human retrosplenial complex during full-body rotation. bioRxiv, 417972.

Groppe, D. M., Makeig, S. and Kutas, M. (2009) Identifying reliable independent components via split-half comparisons. NeuroImage, 45, 1199–1211.

Hyvärinen, A., Karhunen, J. and Oja, E. (2001) Independent Component Analysis. New York: John Wiley & Sons.

Hyvärinen, A. and Oja, E. (2013) Independent Component Analysis: Algorithms and Applications. Neural Networks, 56, 963–976.

Jungnickel, E., Gehrke, L., Klug, M. and Gramann, K. (2018) MoBI-Mobile Brain/Body Imaging. In Neuroergonomics: The Brain at Work and in Everyday Life (eds. H. Ayaz and F. Dehais), chap. 10, 59–63. London: Elsevier, 1st edn.

Ladouce, S., Donaldson, D. I., Dudchenko, P. A. and Ietswaart, M. (2017) Understanding Minds in Real-World Environments: Toward a Mobile Cognition Approach. Frontiers in Human Neuroscience, 10, 1–14.

Ladouce, S., (2019) Mobile EEG identifies the re-allocation of attention during real-world activity. Scientific Reports, 9, 1–10.

Leutheuser, H., Gabsteiger, F., Hebenstreit, F., Reis, P., Lochmann, M. and Eskofier, B. (2013) Comparison of the AMICA and the InfoMax algorithm for the reduction of electromyogenic artifacts in EEG data. Proceedings of the Annual International Conference of the IEEE Engineering in Medicine and Biology Society, EMBS, 6804–6807.

Makeig, S., Gramann, K., Jung, T. P., Sejnowski, T. J. and Poizner, H. (2009) Linking brain, mind and behavior. International Journal of Psychophysiology, 73, 95–100.

Makeig, S., Jung, T. P., Bell, A. J., Ghahremani, D. and Sejnowski, T. J. (1997) Blind separation of auditory event-related brain responses into independent components. Proceedings of the National Academy of Sciences of the United States of America, 94, 10979–10984.

Palmer, J. A., Kreutz-delgado, K. and Makeig, S. (2011) AMICA : An Adaptive Mixture of Independent Component Analyzers with Shared Components. 1–15.

Pion-Tonachini, L., Kreutz-Delgado, K. and Makeig, S. (2019) ICLabel: An automated electroencephalographic independent component classifier, dataset, and website. NeuroImage, 198, 181–197.

Protzak, J. and Gramann, K. (2018) Investigating established EEG parameter during real-world driving. Frontiers in Psychology, 9, 1–11.

Richer, N., Downey, R. J., Nordin, A. D., Hairston, W. D. and Ferris, D. P. (2019) Adding neck muscle activity to a head phantom device to validate mobile EEG muscle and motion artifact removal. 2019 9th International IEEE/EMBS Conference on Neural Engineering (NER), 275–278.

Wascher, E., Heppner, H. and Hoffmann, S. (2014) Towards the measurement of event-related EEG activity in real-life working environments. International Journal of Psychophysiology, 91, 3–9.

Widmann, A., Schröger, E. and Maess, B. (2015) Digital filter design for electrophysiological data – a practical approach. Journal of Neuroscience Methods, 250, 34–46.

Winkler, I., Debener, S., Muller, K. R. and Tangermann, M. (2015) On the influence of high-pass filtering on ICA-based artifact reduction in EEG-ERP. Proceedings of the Annual International Conference of the IEEE Engineering in Medicine and Biology Society, EMBS, 2015-Novem, 4101–4105.

Wunderlich, A. and Gramann, K. (2018) Electrocortical evidence for long-term incidental spatial learning through modified navigation instructions. In Spatial Cognition XI (eds. S. Creem-Regehr, J. Schöning and A. Klippel), 261–278. Cham: Springer International Publishing.

Zakeri, Z., Assecondi, S., Bagshaw, A. and Arvanitis, T. (2014) Influence of Signal Preprocessing on ICA-Based EEG Decomposition. IFMBE Proceedings, 41, 563–566.

